# Continuous Monitoring of Glucose and Oxygen using an Insertable Biomaterial-based Multianalyte Barcode Sensor

**DOI:** 10.1101/2024.07.21.604502

**Authors:** Ridhi Pradhan, David Chimene, Brian S. Ko, Artem Goncharov, Aydogan Ozcan, Michael J. McShane

## Abstract

Chronic diseases including diabetes, cardiovascular diseases, and microvascular complications contribute significantly to global morbidity and mortality. Multiplexing technologies offer a promising approach for the simultaneous detection and management of comorbidities, providing comprehensive disease insights. In this work, we describe a miniaturized optical “barcode” sensor with high biocompatibility for continuous monitoring of glucose and oxygen. This enzymatic sensor relies on oxygen consumption in proportion to local glucose levels and the phosphorescence reporting of tissue oxygen with a lifetime-based probe. The sensor was designed to operate in a tissue environment with low levels of dissolved oxygen. The barcode sensor consists of a poly(ethylene) glycol diacrylate (PEGDA) hydrogel with four discrete compartments separately filled with glucose or oxygen-sensing phosphorescent microparticles. We evaluated the response of the barcode hydrogels to fluctuating glucose levels over the physiological range under low oxygen conditions, demonstrating controlled tuning of dynamic range and sensitivity. Moreover, the barcode sensor exhibited remarkable storage stability over 12 weeks, along with full reversibility and excellent reproducibility (∼6% variability in phosphorescence lifetime). Electron beam sterilization had a negligible impact on the glucose response of the barcode sensors. Furthermore, our investigation revealed minimal phosphorescence lifetime changes in oxygen compartments while exhibiting increased lifetime in glucose-responsive compartments when subjected to alternating glucose concentrations (0 and 200 mg/dL), showcasing the sensor’s multianalyte sensing capabilities without crosstalk between compartments. Additionally, evaluation of tissue response to sensors inserted in pigs revealed appropriate biocompatibility of the barcodes.

---

Chronic diseases are long-lasting health-related problems affecting the quality of life of many people and causing morbidity and mortality worldwide. Chronic disease causes nearly 41 million deaths every year, equivalent to 74% of all deaths globally^1^. According to the World Health Organization (WHO), chronic diseases such as cardiovascular disease (CVD), cancer, stroke, chronic obstructive pulmonary disease, and diabetes are the leading causes of death and disability in the United States. In many cases, patients experience comorbidities, such as diabetes and chronic renal failure, or diabetes and microvascular diseases^2^.

Diabetes is the most common endocrine disorder and a leading cause of mortality worldwide. According to the WHO, in 2019 diabetes mellitus was responsible for 1.5 million deaths. Additionally, 460,000 kidney disease deaths and 20% of cardiovascular deaths were caused by diabetes^1^. Continuous glucose monitoring (CGM) provides real-time glycemic monitoring to improve diabetes management and health outcomes. In addition to glucose monitoring, regional tissue oxygen is a key interest in detecting and managing microvascular disease in diabetic patients^3, 4^. The reduced regional oxygen levels in diabetic patients can contribute to the progression of complications associated with microvascular diseases, including nephropathy, retinopathy, and albuminuria. Consequently, measuring regional tissue oxygen emerges as a crucial aspect in identifying microvascular diseases in diabetic patients^4^. Hence, employing a multi-analyte sensor capable of assessing tissue oxygen levels, alongside continuous glucose monitoring, could serve as an invaluable diagnostic tool for preventing microvascular diseases and effectively managing diabetes in affected individuals.

The commercially available electrochemical CGMs such as Abbott Libre, Dexcom, and Medtronic Guardian that are partially implanted pose constant sensor tissue friction, shortening the longevity of these sensors to 3-14 days^5, 6^. Alternatively, optical measurements including spectroscopy (NIR, Raman), and fluorescence have shown high potential in long-term implantable continuous glucose sensing^7, 8^. Eversense, an optical-based CGM (measuring 3.5 × 18.3 mm) has been developed by Senseonics, which is implanted into the subcutaneous tissue of the upper arm through a surgical procedure using local anesthesia. Currently, only this sensor presents a longer lifespan (180 days) compared to electrochemical CGM sensors. Nevertheless, challenges related to invasiveness and biocompatibility (foreign body reaction -FBR) persist. FBR hampers metabolite diffusion and degrades optical chemistry, ultimately causing inaccuracies in the sensor’s readings. The intensity and nature of the inflammation cascade are intricately linked to the initial inflammatory response, particularly the adsorption of biomolecules, as well as the characteristics of the foreign body such as size, shape, and physical and chemical properties^9, 10^. Therefore, miniaturizing implantable CGMs is crucial to ensure less invasive insertion and minimize the risk of potential immune reactions against the sensor.

A continuous monitoring system offers a prolonged tracking of physiological parameters and plays a critical role in chronic disease management. Continuous monitoring devices can detect abrupt or unforeseen fluctuations between discrete measurement times, enabling early intervention, personalized care, and reduced hospitalizations by identifying changes in real-time^5^. The notable success of continuous glucose monitoring (CGM) to improve the therapeutic outcome of patients over the past decades has sparked a growing interest among researchers and clinicians in the continuous sensing of additional biomarkers8. However, in some cases, the existence of comorbidities may only be identified by monitoring the concentration of multiple biomarkers. Additionally, relying solely on the detection of a single biomarker often needs to be revised for clinical diagnosis or monitoring disease progression. Multiplexing enables simultaneous detection of multiple analytes which can significantly improve the management of comorbidities by providing more information about the disease(s), and its/their status^11, 12^. Multianalyte sensors also have the advantage of providing a shorter overall analysis time for multiple analytes within the sample volume offering a cost-effective and rapid analysis over biosensing of individual analytes.

As an alternative to electrochemical sensors, some recent papers have reported on insertable optical sensors that do not require implanted electronics; these typically rely on phosphorescent oxygen-sensitive luminescent dye embedded into hydrogel matrices^13-15^. To make these materials responsive to other analytes, oxidoreductase enzymes are also included to enable measurement of, for example, glucose, oxygen, and other analytes^16-18^. In these systems, a luminescent dye/ phosphor is collisional quenched by oxygen, reducing its phosphorescence lifetime; thus, the lifetime is proportional to the local oxygen concentration. The biosensing assays indirectly respond to a target analyte by depleting local oxygen, mediated by specific oxidoreductase enzymes. This depletion alters the phosphorescence lifetime in proportion to the analyte concentration: the reduction of oxygen in the system increases the phosphorescence lifetime, which is then used to estimate the concentration of the target analyte using a known relationship determined by pre-calibration.

We have recently reported a multi-compartment poly(ethylene)glycol diacrylate (PEG-DA) miniaturized implantable hydrogel called the “barcode” which has dimensions comparable to a rice grain and can be easily inserted into subcutaneous tissue using a 16-gauge needle^19^. These barcode sensors encapsulated discrete assays based on phosphorescent Pd(II) metalloporphyrin dyes for continuous monitoring of glucose and oxygen with minimal crosstalk. The glucose-sensing microparticles and oxygen-sensing microparticles consisted of two different porphyrin dyes. The glucose-responsive microparticles consisted of Pd (II) meso-Tetra (sulfophenyl) Tetrabenzoporphyrin Sodium Salt (HULK) and the oxygen sensing microparticles consisted of [Pd-meso-tetra(4-carboxyphenyl) porphyrin (PdP). The HULK has a red excitation wavelength of 630 nm which falls within the optical window. However, PdP has an excitation wavelength near 530 nm, which can cause difficulty in excitation in vivo due to the absorption and scattering of light. In addition, it is notable that those initial proof-of-concept experiments for these barcode sensors were performed at oxygen levels much higher than those of interstitial tissue oxygen7. Hence, these sensors quickly saturated at relatively low analyte concentrations when used in typical low oxygen concentrations found in tissue (∼30-40uM)^4^.

This work aimed to advance the barcode sensor for improved sensitivity and stability by optimizing the design for performance at lower interstitial oxygen conditions. To accomplish this, we replaced the HULK porphyrin dye with oxygen-sensitive ethyl cellulose nanoparticles (ECNP) encapsulating a palladium (II) meso-tetra(4-carboxyphenyl)tetrabenzo-porphyrin) dye (PdBP). ECNPs have proved to be more sensitive compared to the free form of the dye^20^. Additionally, to tune the sensitivity of these sensors under lower oxygen conditions, the glucose-sensing microparticles were further modified with crosslinked diffusion-limiting polyelectrolyte coatings to control the glucose permeation more precisely into the sensing domains. Furthermore, the stability, reproducibility, effects of sterilization, reversibility, and biocompatibility of the barcode sensors were assessed.

## EXPERIMENTAL SECTION

### Reagents and Materials

Alginate (Cat No. A2158, 75-100 kDa), ethyl cellulose (EC, Cat No. 200697, 48% ethoxy), TRIS base (Cat No. T1503), [2-(Methacryloyloxy) ethyl]trimethylammonium chloride (TMA, Cat. No. 408107, 80% in water), catalase (CAT, from bovine liver, Cat. No. C9322), calcium chloride (Cat No. 22350), pluronic F 68 (PF68, Cat. No. P1300), poly (allylamine hydrochloride) (PAH, Cat No.71550-12-4, 17.5 kDa), poly(sodium-4-styrenesulfonate) (PSS, Cat No. 25704-18-1, 70 kDa), and tetrahydrofuran (THF, Cat No. 401757) were purchased from Sigma-Aldrich, Inc., St. Louis, MO, USA. Palladium (II) meso-tetra(4-carboxyphenyl)tetrabenzo-porphyrin) (PdBP, Cat No. T13343) was purchased from Frontier Specialty Chemicals, Logan, UT, USA. Iso-octane (Cat No. 94701) was purchased from Avantor performance materials, LLC, Randor, PA, USA. Glucose oxidase (GOx, Cat No. 9001-37-0, Activity-76.8 unit/mg) from Aspergillus niger was purchased from Tokyo Chemical Industries. Co (Tokyo, Japan). Poly (ethylene glycol) diacrylate (PEGDA, average Mw ∼ 3.4 kDa) was purchased from Alfa Aesar (Haverhill, MA, USA). 2,2-dimethoxy-2-phenyl acetophenone (C6H5COC(OCH3)2C6H5,> 99%), 1-vinyl-2-pyrrolidinone (C6H9NO, >99%) were obtained from Sigma-Aldrich (St. Louis, MO, USA).

### Synthesis of Nanoparticles Containing Oxygen-Sensitive Phosphors

The oxygen-sensitive nanoparticles were fabricated using the previously reported nano-emulsion method^20^. Briefly, 100mg of EC was dissolved in 5 ml of THF overnight using a magnetic stirrer. Then, 2 mg of PdBP was dissolved into the above solution of EC and THF using sonication for 30 minutes. The above THF solution with dye and polymer was filtered through a 0.2 μ PTFE syringe filter and stored in a vial. In a separate 50 mL centrifuge tube, 100 mg of surfactant (PF 68) was dissolved in 20 mL of nanopore water using sonication and filtered through a 0.2 μ PTFE syringe filter. Next, the surfactant solution was sonicated using a sonication probe, and the THF solution with dye and polymer was slowly injected into the aqueous solution of surfactant within 30 of the 2-minute sonication process. Finally, the suspension of NPs was filtered through a 100 μ nylon filter to remove larger precipitates, and the volume was reduced to 3 mL using centrifuge filtration. Then, the NP suspension was washed with 15mL nano pure water to remove excess surfactant. These NPs were stored in a vial of 16.6 mg/mL in the fridge (4ºC) for further use.

### Synthesis of Sensing Microparticles with Nanofilms

Alginate microparticles encapsulating the oxygen-sensitive ECNPs, glucose oxidase (GOx), and catalase (Cat) were synthesized using an emulsion technique. A homogenous mixture of 3.75mL of 4% w/v aqueous solution of sodium alginate and 1.25 mL (5 mg NPs) of nanoparticle suspension was obtained by nutating the mixture for 30 minutes. In a separate tube, 58.5 mg of GOx and 54.9 mg of Cat were dissolved in 2.5 mL of 50 mM TRIS buffer (pH 7.2) by gentle nutation. Then, the alginate mixture and the enzyme mixture were mixed to prepare a precursor solution. This mixture was added dropwise and emulsified for 2 minutes in a solution containing 10.8 mL isooctane with 322 μL SPAN 85 using a homogenizer operating at 8000 rpm. Next, 1.5mL of isooctane with 175 μL TWEEN 85 was added to the above mixture and stirred with the same speed for 15 seconds. During the last 50 seconds of the emulsification, 4 mL of 10 w/v% CaCl2 solution was added to allow external gelation of the alginate microparticles. The emulsion was then transferred to a round bottom flask and gently stirred in a magnetic stirrer for 20 minutes. The microparticles were centrifuged at 2000 g for two minutes and washed with deionized water two times. Next, the surfaces of the microparticles were modified by applying ultrathin nanofilms of polyelectrolytes using the electrostatic layer-by-layer (LbL) assembly. Figure 1 illustrates the fabrication of sensing alginate microparticles bound with polyelectrolyte nanofilms.

**Figure 1:**
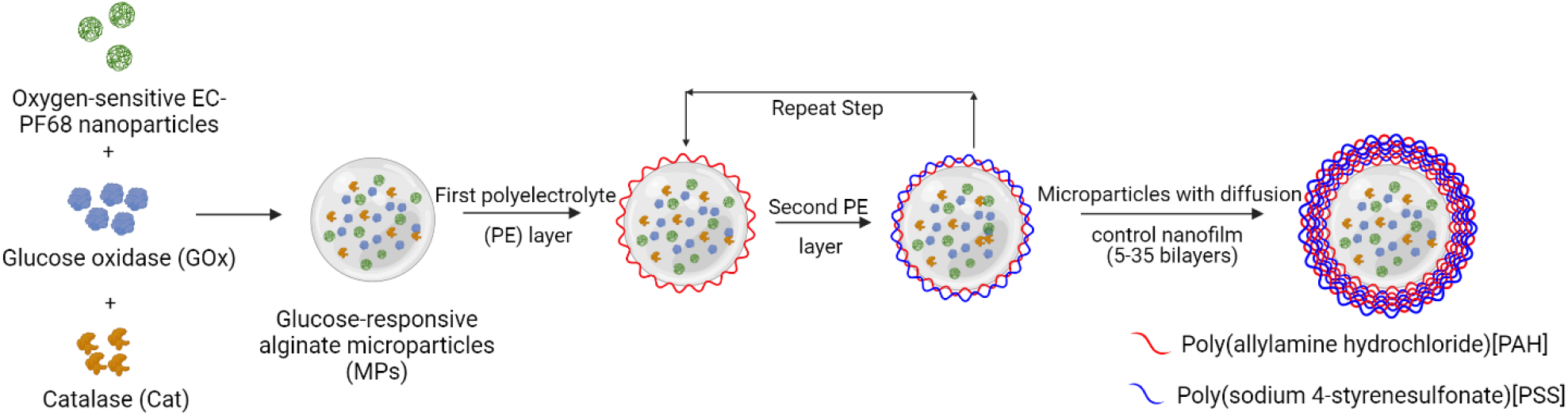
Illustration of alginate microparticle encapsulating oxygen-sensitive ethyl cellulose nanoparticles (ECNP), glucose oxidase (GOx), and catalase (Cat) coated with polyelectrolyte multilayer for glucose diffusion control. Multilayer nanofilms were deposited by layer-by-layer sequential electrostatic adsorption method.

The oxygen-sensing microparticles were synthesized using the same method used for the glucose-sensing microparticles excluding the enzymes.

### Layer-by-Layer Deposition

Polyelectrolyte nanofilms were deposited on the alginate microparticles by dispersing the pallet of the microparticles in 2ml of PAH (pH 8) followed by centrifugation and the supernatant was discarded. Then the pallet was resuspended in PAH wash solution (10mM TRIS buffer with pH 8), centrifuged and the supernatant was discarded. The same procedure was followed for PSS (pH 7.2) and PSS wash solution (10 mM TRIS buffer with pH 7.2). This created one bilayer of polyelectrolytes PAH and PSS. The microparticles for oxygen sensing consisted of 5 bilayers. The microparticles for glucose sensing consisted of a total of 5, 10, 15, 20, 25, 30 and 35 bilayers. Further, covalent crosslinking of the amine group of PAH of the glucose-sensing microparticles was also performed by mixing 8.8 mg of PEM-coated microparticles with 2 mL of 0.1 M glutaraldehyde and stirring for 30 minutes. The microparticles were then suspended in a 10mM TRIS buffer and stored at 4ºC for future use.

The assembly of polyelectrolyte layers on alginate microspheres was monitored by electrophoretic mobility measurements (Zeta potential, Malvern ZetaSizer Namo ZS). Cellometer Mini (Nexcelom) was used to determine the size of alginate microparticles after LbL depositions. The stock microparticle solution was diluted 1:20 in the TRIS buffer. Then, 20 μL of the diluted solution was dispensed onto the slide for analysis.

### Fabrication of Barcode Hydrogel Sensor

To create a discrete compartment barcode sensor, a previously established soft lithography process was used^19^. A PDMS top and bottom master mold were fabricated by replica molding from a 3D-printed master mold. The PDMS, the precursor, and the curing agent were mixed at a ratio of 10: 1, poured into the printed master mold and cured at 60°C under vacuum for 2 hours. The hydrogel solution was prepared by mixing 20% (w/v) PEGDA and 2% (v/v) of the photo-initiator solution. The hydrogel precursor solution was dispensed into the bottom master mold, and the top master mold was aligned. The hydrogel was crosslinked under a UV lamp (360 nm, 10-15 mW cm^−2^) by 5 min exposure. The hydrogel case was then peeled from the PDMS mold and rinsed in deionized water (DI water).

The sensing assay (8.8mg/90μL) was mixed with the hydrogel precursor in a ratio of 3:1. For the glucose sensor termed as “glucose barcode”, glucose sensing microparticles were mixed with the hydrogel precursor, and the “oxygen barcode” had only oxygen sensing microparticles. Then, 0.64 μL of the mixture was pipetted into 2 compartments and polymerized under UV for 5 minutes. For the multianalyte barcode containing glucose and oxygen sensing assay, two alternate compartments were filled with a mixture of hydrogel precursor and glucose sensing microparticles, and the remaining compartments were each filled with a mixture of hydrogel precursor and oxygen sensing microparticles. After polymerization, the barcode sensors were rinsed in DI water and stored in a TRIS buffer at 4°C. Figure 2 illustrates the fabrication process of barcode hydrogel sensors.

**Figure 2:**
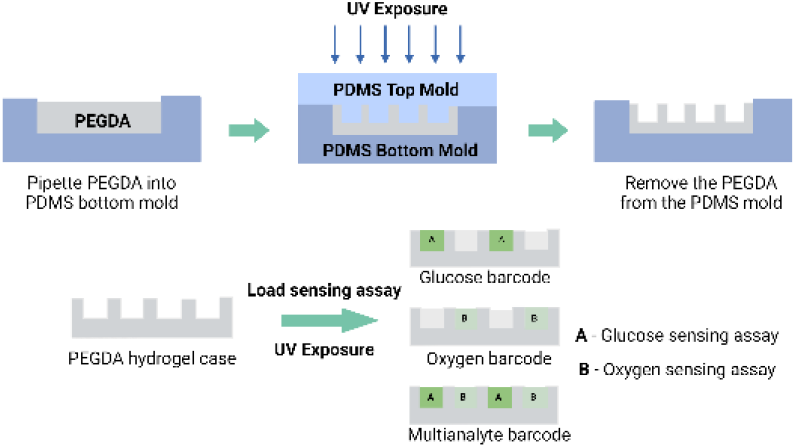
Illustration of the fabrication process of barcode hydrogel sensor using soft lithography.

### Characterization of Oxygen and Glucose Response

The barcode hydrogels (n=4) were immobilized in a previously described custom-built flow cell consisting of an acrylic sheet that holds up to four samples inside and attaches optical readers from outside^21^. Oxygen challenges were performed by exposing the sensors to varying dissolved oxygen concentrations inside an incubator at 37°C. The dissolved oxygen concentration of 0-257.9 μM was achieved by mixing air and nitrogen in a defined ratio between 0 to 21% using a digitally-controlled mass flow controller (MKS Instruments PR 4000B). The glucose challenges were performed by exposing the sensors to varying physiologically-relevant glucose concentrations (0-400mg/dL) in an incubator (37°C), while holding oxygen at a fixed concentration of ∼40 μM representative of expected tissue levels. The glucose concentration was modulated using peristaltic pumps (Masterflex, 7550-50) by mixing feeds drawn from reservoirs containing 0 and 400mg/dL of glucose stock solution in 10mM TRIS with 10mM CaCl2. The pump speeds were controlled by a custom LABVIEW program (figure S1) to precisely mix the solutions to achieve glucose levels over the full range of concentrations. The oxygen concentration was regulated using a vacuum degassing chamber (9000-1118) and vacuum pump (9000-1472-Systec)^21^. All lifetime readings were recorded using a custom multichannel time-domain phosphorescence lifetime measurement system with excitation of 630 nm and 800 nm emission^22^.

The multiplexing capabilities of the barcode hydrogel sensors were further tested using a custom phosphorescence lifetime imager (PLI), which can spatially resolve responses from different compartments of the barcode devices^23^. The PLI reader acquired time-lapse phosphorescence images of decaying sensor emissions, which were then processed to obtain phosphorescence intensity and lifetime-coded images of the entire field of view. The lifetime responses of hydrogel sensor compartments were further calculated by averaging pixels within rectangular masks overlaid with each compartment, yielding four signals per device (two glucose-sensitive compartments and two oxygen-sensitive compartments). Final lifetime values for each analyte were obtained by averaging responses from alike compartments^23^.

The limit of detection (LOD) was estimated from a linear regression model by calculating the glucose concentration corresponding to the phosphorescence lifetime at 0 mg/dL glucose plus three times the standard deviation of the lifetime signal at that analyte concentration. Similarly, maximum differentiable glucose concentration (MDGC) was estimated by calculating glucose concentration corresponding to the phosphorescence lifetime at 400 mg/dL glucose plus three times the standard deviation of the lifetime signal at that analyte concentration. The dynamic range was calculated as R= MDGC – LOD, and sensitivity was calculated by dividing the difference in phosphorescence lifetime values at LOD and MDGC by the dynamic range.

### Assessment of Storage and Operational/Enzyme Stability

Barcodes were incubated in two different conditions to evaluate their long-term stability under relevant circumstances. For evaluating the storage stability, samples were stored in a 10mM TRIS buffer containing no glucose at 4°C. The lifetime responses from these sensors were assessed at the beginning, 4, and 12 weeks. In the second condition, the operational/enzyme stability of the barcode sensors was evaluated by incubating samples in physiological solutions of 10mM phosphate-buffered saline (PBS) containing 100 mg/dL glucose at 37°C for 8 weeks. Over the duration of storage, the buffer containing the samples was tightly sealed, stirred, and changed every 4 days. The enzymatic activity of glucose oxidase was monitored using a colorimetric assay that relies on the oxidation of *o*-dianisidine within a peroxidase-coupled system. To conduct the assay, a reaction cocktail consisted of a cocktail of glucose, *o*-dianisidine, and horseradish peroxidase (HRP) in sodium acetate, prepared according to an established protocol^24^. The sensors were placed into a 96-well plate, 200 μL of the reaction cocktail was dispensed into each well, and absorbance at 492 nm was monitored in one-minute kinetic interval cycles for 30 minutes using a plate reader (Biotek Cytation 5). The experiment was repeated for the samples after fabrication and biweekly upon storage in physiological conditions. The change in absorbance was plotted over time, and the slopes of the linear portion of the curves were used to determine the comparative apparent activity among the samples at different time points.

### *In Vivo* Insertion

To better understand the long-term biocompatibility and performance of the barcode samples under real-world conditions, barcode sensors were inserted under the skin of two female pigs. Female piglets were used for ease of handling and so that the pigs could continue to live together after puberty, which occurs roughly at 6 months old. As a production breed, Yorkshire cross pigs are known for robust health and gregarious nature, making them well suited to an extended study. The pigs lived together in semi-enclosed housing, with access to toys and enrichment, including a wading pool and fans, and heat lamps and hay beds as appropriate for the season. This animal study was IACUC-approved under AUP# IACUC 2021-0066 Reference Number: 137661.

The barcode sensors were inserted into the pigs at different time points: at 3 months old in Pig 1, and at 7.5 months old in Pig 2. Accordingly, the barcode sensors were evaluated over 7 months in Pig 1, and 3 months in Pig 2. During insertion, pigs were anesthetized using isoflurane to minimize stress and intubated for safety. The animals were placed on a heating pad and under a blanket where possible to maintain body temperature. Insertion sites were trimmed and shaved, then cleaned with detergent and isopropyl alcohol. A small marking was tattooed around each insertion site because the implants cannot be visually identified otherwise. The sensors were loaded into 16-gauge needles, and inserted subcutaneously to a target depth of 2 mm. A steel dowel rod was used to hold the sensor in place as the needle and dowel rod were withdrawn. After implantation, the pigs were kept under observation until anesthesia had worn off, then returned to her sister in the main pen.

After the experiment, pigs were humanely euthanized by veterinary staff. The fresh hides containing the sensors were then skinned off and stored in 10% neutral buffered formalin for at least 24 hours. The hides were then sectioned off, using the tattoos and lifetime readings to precisely locate each sensor. Roughly one square inch of hide surrounding the strongest sensor reading was then removed, serially sectioned, paraffin-embedded, and stained with hematoxylin & eosin. Independent veterinary pathologists then examined the slides for device presence, device state, skin zone, host interface, and healing response.

### Statistical Analysis

All data are expressed as mean ± one standard deviation (SD). One-way analysis of variance tests was performed in Origin Pro for comparison between groups. A p-value >0.05 was considered statistically significant.

## RESULTS AND DISCUSSION

### Characterization of Alginate Microparticles and Barcode Hydrogels

The size and surface charge of alginate microparticles with different LBLs were characterized using the cellometer and zeta potential analyzer respectively, to ensure proper deposition of polyelectrolyte layers. The diameter of microparticles coated with 5-35 bilayers of PAH/PSS ranged from 10-18μm respectively (Figure 3a). The size of the microparticles increased slightly with the increments of the polyelectrolyte bilayers. The zeta potential results demonstrated the reversal of the surface charge of the microparticles at each step from –20 mV for PSS to +40 mV for PAH (Figure 3b). The increasing size and reversal charge indicate the deposition of PAH/PSS layers on the alginate microparticles^25^.

**Figure 3.**
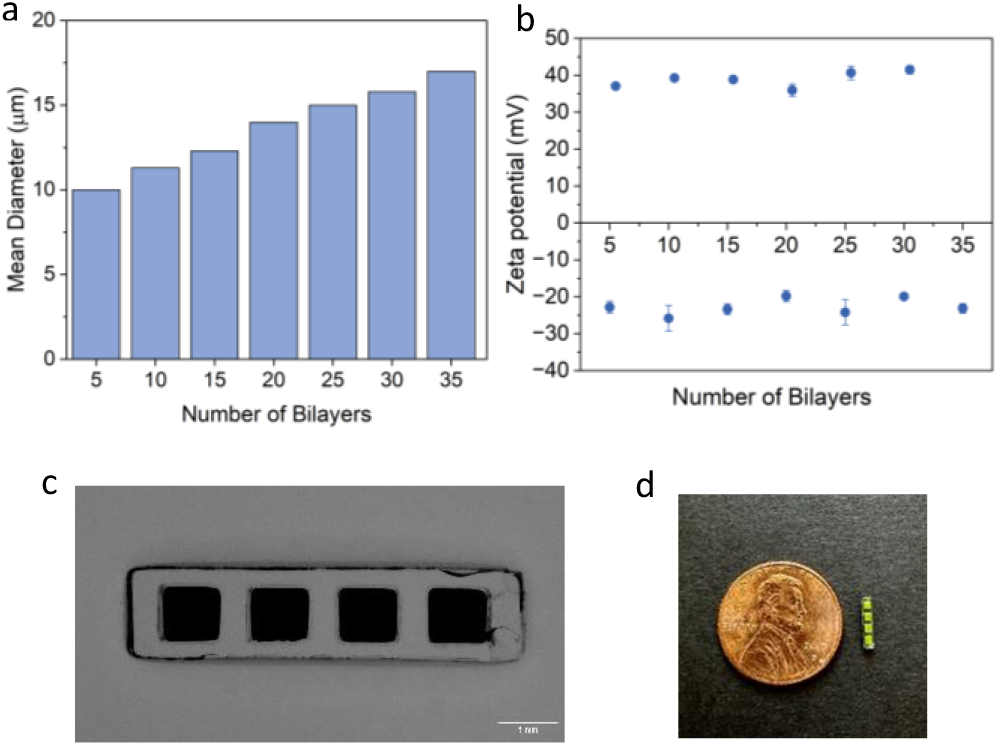
(a) Average diameter of synthesized alginate microparticles bounded with nanofilms of different number of bilayers. (b) ζ-Potential measurements for deposition of polyelectrolyte nanofilms on alginate microparticles. (c) Microscopic image of barcode hydrogel sensor, Scale bar=1mm. (d) Image of barcode hydrogel compared to a penny.

The fabricated barcode hydrogel had a one-by-four linear array of hollow cuboid structures with overall dimensions of 6.5 mm in length, 1.2 mm in width, and 1 mm in height. The individual compartments were each 1 mm long, 0.8mm wide, and 0.8mm tall and the compartments were separated by 0.5 mm of hydrogel. The stereotaxic microscopy images of the barcode hydrogel (Figure 3c) demonstrate the filled compartments, which appear darker than the hydrogel casing.

### Barcode Oxygen and Glucose Response

The lifetime response of the glucose and oxygen barcode sensors to the changes in the oxygen concentrations was measured. The linear Stern Volmer relationship: τ_0_/τ = 1 + K_SV_[O_2_] was used to evaluate the oxygen response. The oxygen-sensitive compartments exhibited a pronounced response to changes in the oxygen concentration from 0% to 21% (0-257.9μM), as expected (Figure 4, S2), with a phosphorescence lifetime range of 61-272 μs and a Stern Volmer constant (K_SV_) value of 0.16 %O_2_^1^. The response of the glucose-sensitive compartments were also evaluated for response to the changing oxygen concentration, from which it was determined that was no significant difference in the K_SV_ (summarized in Table S1) between these compartments and the oxygen sensors; further, there was no significant difference among the particles with different bilayers of nanofilms whether crosslinked or not, indicating the nanofilms did not substantially affect the kinetics of oxygen diffusion (Figure 5).

**Figure 4.**
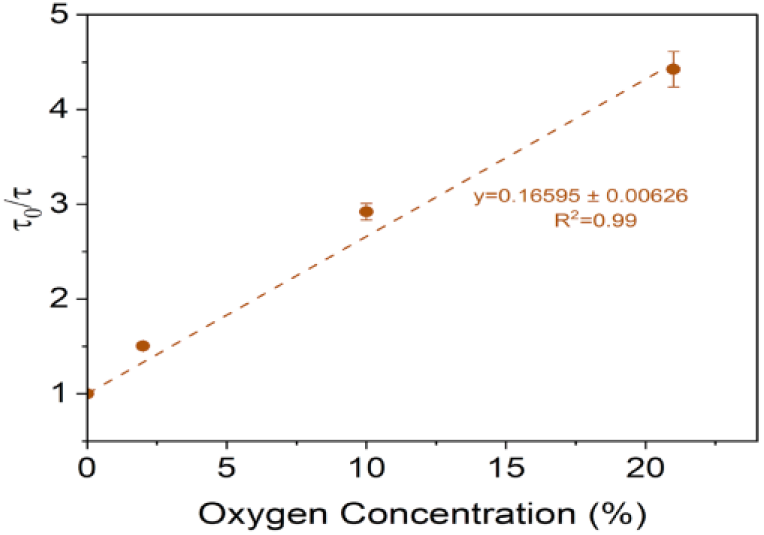
Stern-Volmer plot for oxygen barcode hydrogel sensor under different oxygen concentrations. Error bars represent the SD from the mean for N=4 different samples.

**Figure 5.**
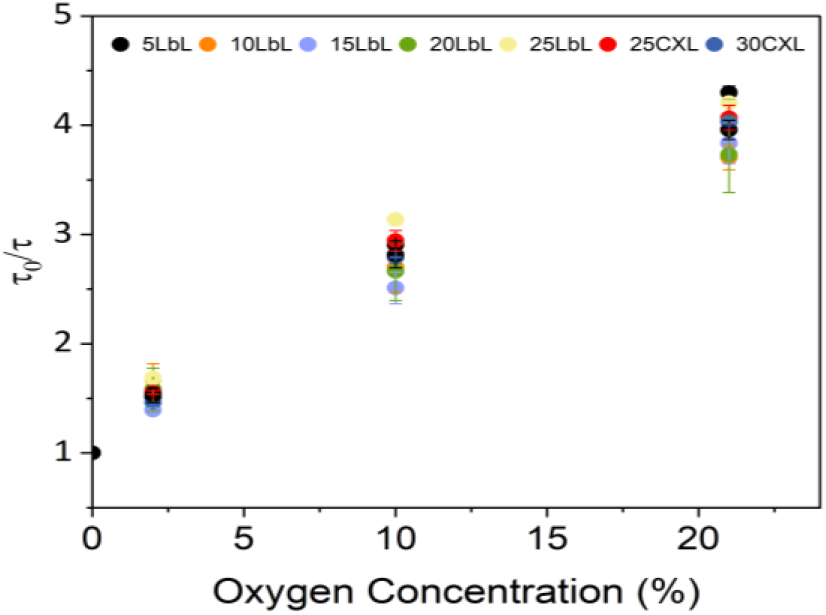
Stern-Volmer plots for glucose barcodes packed with alginate microparticles bound by varying nanofilm coatings under different oxygen concentrations. Error bars represent the SD from the mean for N=4 different samples.

The glucose barcode hydrogels were subjected to testing at physiologically-relevant glucose concentrations (0-400 mg/dL) under a hypoxic oxygen environment (35-40 μM). When the number of bilayers used to construct the nanofilm coatings was increased from n=5 to n=25, the dynamic range of response to glucose exhibited an increase of ∼145%. The non-crosslinked microparticles with the most bilayers (n=25) yielded a maximum detectable concentration of 133 mg/dL with a sensitivity of 0.37 us-dl/mg. The expansion of the dynamic range of these sensors is directly related to the increase in the number of bilayers, an effect attributed to a decrease in the flux of glucose into the sensing domains with thicker films, thus decreasing the sensitivity. This result aligns with previous reports on flux-based enzymatic glucose sensors which showed an increase in dynamic range and a simultaneous decrease in sensitivity, with the decrease of glucose diffusion while maintaining the oxygen diffusion rates^26^. Despite having 25 bilayers, when tested in the hypoxic conditions the barcode sensor exhibited sensitivity to glucose levels only in the hypoglycemic range (0-100mg/dL); the lifetime saturated between 225 and 250us and failed to further respond to the hyperglycemic levels as shown in Figure 6a. This suggests that glucose diffusion through the sensing domain remains high, resulting in consumption of the limited oxygen even at lower glucose levels. Hence, to further decrease the diffusion of glucose into the sensing domains and extend the concentration range of the response in hypoxic conditions, the nanofilms were crosslinked using glutaraldehyde. It is known that when the PAH/PSS nanofilms are treated with glutaraldehyde, the amino group of the PAH in the nanofilm readily reacts with the aldehyde group of glutaraldehyde^26^; this has previously been proven to be an effective method to reduce glucose permeation into the microparticles without hindering the diffusion of oxygen^17, 26^. Crosslinking the 25-bilayer nanofilms increased the dynamic range by ∼16% yielding a maximum detectable concentration of 175 mg/dL. Further increasing the crosslinked nanofilms to 35 bilayers further increased the dynamic range by 84% (Figure 6b). The sensor demonstrated linear response (R2>0.99) within the physiological glucose concentrations of 0-400 mg/dL. The lower range of LOD and MDGC of the sensors was 37 mg/dL and 321 mg/dL respectively, with a sensitivity of 0.19. The dynamic range and sensitivity of each sensor type are summarized in Table 1. Hence, considering crucial factors like linear response and sensitivity, MPs with 30 LbLs with glutaraldehyde crosslinked nanofilms were selected for subsequent in vitro analysis.

**Figure 6.**
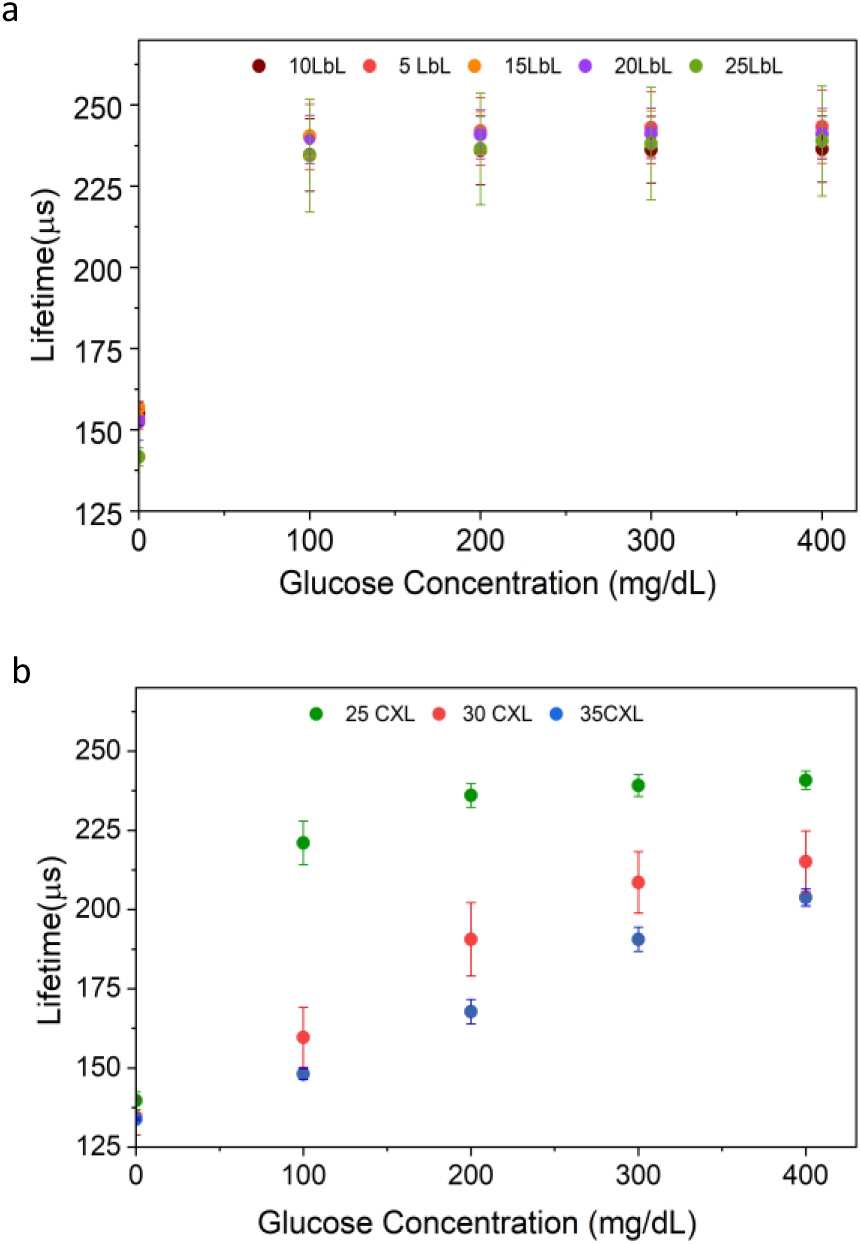
Steady-state lifetime response plots of glucose barcodes (n=4) packed with alginate microparticles with different LbLs, dispersed in barcode hydrogel by challenging the sensors to physiological glucose concentration (0-400 mg/dL) at 40 μM dissolved oxygen concentration and 37°C. (a) Sensors consisted of non-crosslinked nanofilm coatings. (b) Sensors consisted of glutaraldehyde crosslinked (CXL) nanofilm coatings. Error bars represent the SD from the mean for N=4 different samples.

### Sterilization and Reproducibility

Sterilization is a crucial prerequisite for implantable sensors to ensure their safe and effective use in clinical applications. However, enzymatic sensors can be sensitive to sterilization processes, leading to loss of sensor activity. Specifically, using heat and toxic gasses such as ethylene oxide for sterilization of enzymatic sensors can cause irreversible denaturation of the enzymes^27^. As a result, radiation-based sterilization has become a widely adopted approach for enzymatic sensors. Here, the barcode sensors were sterilized using electron beam (e-beam) radiation, which offers advantages in terms of affordability and ease of control when compared to gamma radiation.

The barcode sensors were subjected to e-beam irradiation in a 10mM TRIS buffer with a dose of <25kGy. The lifetime response towards increasing glucose concentration showed only a minor impact, with a lifetime change of ∼6% between non-sterilized and e-beam sterilized barcode sensors. There were no significant differences (p> 0.05) observed at glucose concentrations of 200 mg/dL and 400 mg/dL, as shown in Figure 7a. Previous studies involving enzymatic glucose and lactate sensors sterilized with the same method exhibited functional sensors but with sensitivity reductions up to 60% after radiation^28, 29^. Dang et al observed that when the optical enzymatic sensors were stored in a 5 mM TRIS buffer during radiation, protective effects on the proteins were observed, resulting in an 80% retention of response when exposed to 15kGy^30^. TRIS buffers have radical scavengers that can absorb the free electron energy produced during the e-beam process, thus protecting the enzyme by mitigating the negative effects of e-beam sterilization^31^. This indicates that immersing the barcode sensors in a TRIS buffer during and after e-beam radiation may have minimized radiation-induced damage.

**Figure 7.**
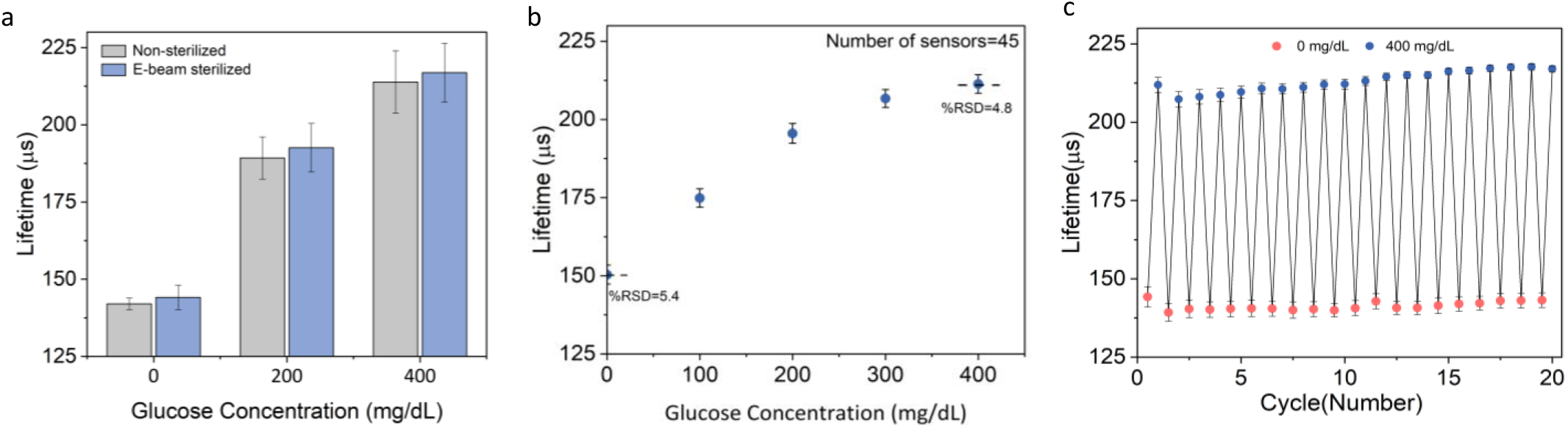
(a) Effect of 20kGy electronic beam (Ebeam) sterilization on barcode glucose sensors (N=8). (b) Reproducibility of the barcode glucose sensors (N=45 different sensors) for detection of glucose ranging from 0-400 mg/dL of glucose with RSD value <6%. (c) Reversibility of barcode sensors (N=4) exposed to 0 and 400 mg/dL of glucose. Error bars represent SD from the mean.

The sensor-to-sensor reproducibility was also examined by measuring the lifetime responses of 45 replicate barcode sensors consisting of sensing alginate microparticles from two different batches. All the sensors were fabricated under similar conditions and exposed to glucose concentrations ranging from 0-400 mg/dL for testing. This yielded an acceptable reproducible response with a relative standard deviation (RSD) in a lifetime measurement of 6% (Figure 7b). To investigate the reversible glucose response, the barcode sensors were exposed to 20 consecutive cycles of 0 and 400 mg/dL of glucose concentrations (Figure 7c). The average lifetime response of the sensor at 0mg/dL glucose concentration was 131±2.6μs while the average lifetime response of the sensor at 400mg/dL glucose concentration was 213±3.2μs. Together these findings showed that the barcode sensors can be fabricated to have a very consistent response, and they exhibit excellent reversibility.

### Storage and Operational/Enzyme Stability

Potential for long-term storage and operational stability are required for the continuous monitoring of metabolites. Over time, enzymatic sensors can degrade due to two factors: enzyme denaturation and hydrogen peroxide poisoning. Hence, both the storage and operational stability of the barcode sensors involving the enzymes were evaluated.

The lifetime response from barcode sensors stored at 4°C showed no significant changes after 12 weeks. The sensors retained 94% of their initial response when exposed to 400 mg/dL of glucose after 12 weeks, as shown in Figure 8a. The relatively high storage stability of the barcode sensors may be due to the immobilization of glucose oxidase within the hydrogel microparticles which has proven to be an effective method of preserving enzyme activity over an extended period^25, 31^. For determining the operational stability of the barcode sensors, both the enzyme activity and lifetime response were evaluated. Compared to the standard GOx enzyme solution (GOx dissolved in TRIS buffer) which could retain only 3% of the activity by 8 weeks, the barcode sensors were observed to retain 80% of their apparent activity over the first four weeks of incubation. However, after 8 weeks of incubation, only 35% of the initial apparent activity was retained (Figure 8a). These findings of enzyme activity loss matched with the observed plateau at less than 200 μs in the lifetime response at 200 mg/dL of glucose after 5 weeks of incubation, indicating that the system had become enzyme-limited (Figure S4). Under these conditions, a fraction of the enzyme activity loss may also be due to hydrogen poisoning leading to enzyme denaturation. However, the preserved enzyme stability during operation should be helped by the co-immobilization of CAT which has been reported to mitigate the effect of hydrogen peroxide produced during glucose catalysis^32^. Additionally, the incorporation of PAH/PSS nanofilm coating has been reported to sustain the GOx activity encapsulated in alginate microspheres^25^.

**Figure 8.**
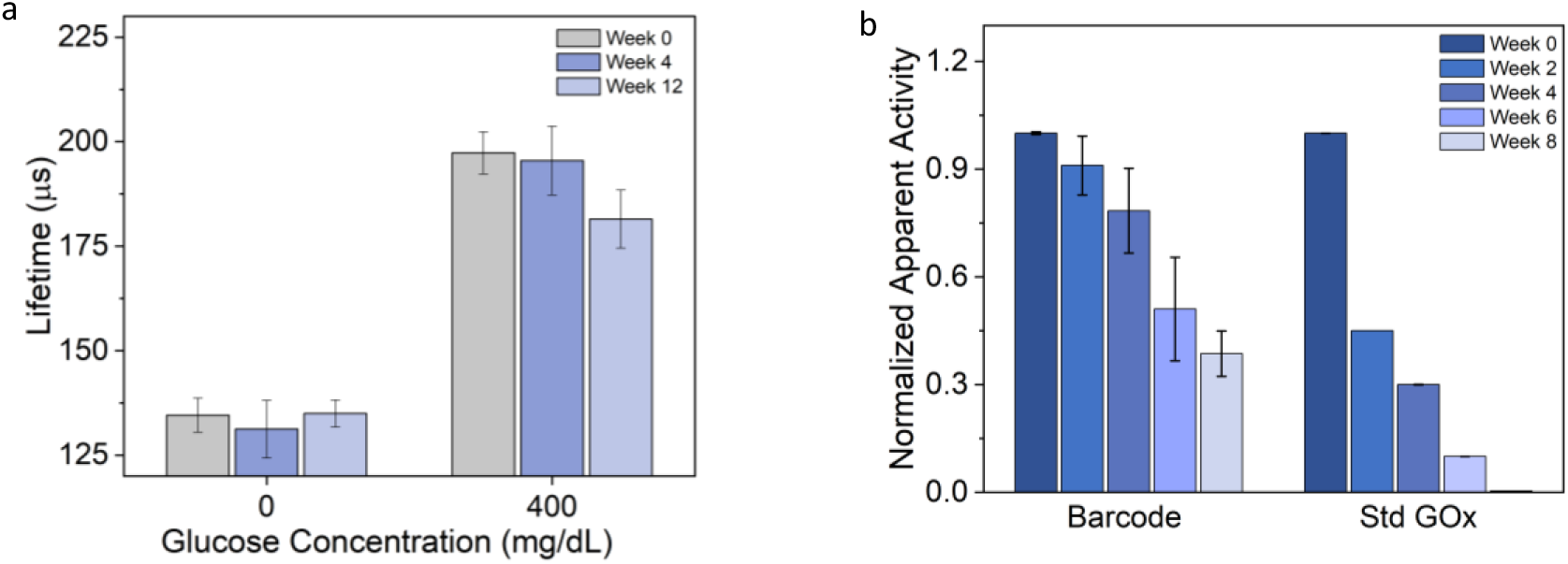
(a) Storage stability of barcode glucose sensors over 12 weeks under storage condition of 4°C in 10mM TRIS buffer. (b) Enzyme stability of barcode glucose sensors compared to dissolved glucose oxidase over 8 weeks under storage condition of 37°C in PBS with 100 mg/dL of glucose. Error bars represent the SD from the mean for N=4 different samples.

### Multi-analyte System Response

A crucial aspect of multi-analyte detection lies in managing the cross-sensitivity between assays, as well as signal crosstalk. Since the phosphorescence lifetime changes from the glucose and oxygen assays are both sensitive to oxygen depletion, the diffusion of oxygen molecules between the neighboring compartments could potentially introduce crosstalk in the barcode sensors if too physically close. For this work, the distance between each sensing compartment was 0.5 mm, which was found in previous experiments to avoid crosstalk^19^.

To assess crosstalk between compartments, the multianalyte barcode sensors were subjected to cycles of physiologically relevant glucose concentrations (0-200 mg/dL) at a constant oxygen concentration of 40uM at 37°C. The barcodes were left to reach a steady state for 20 minutes between the subsequent glucose changes. The multiplexed lifetime measurements were obtained using the phosphorescence imaging (PLI) reader, which acquired a series of phosphorescence intensity images of the entire multi-analyte sensor and used these images to derive the lifetime responses of glucose-sensitive and oxygen-sensitive compartments.

It was found from these data that the baseline lifetime in the absence of glucose was comparable for both types of compartments, as expected. Further, the phosphorescence lifetime in glucose-sensitive compartments increased and decreased by 20% when glucose concentrations oscillated between 0 to 200 mg/dL across cycles. Conversely, the glucose-insensitive (oxygen-sensitive) compartments exhibited minimal phosphorescence lifetime variation of only 0.1% in response to changes in glucose concentration (Figure 9). The slight variations in the oxygen compartments following the glucose modulation indicate negligible crosstalk between the sensing compartments.

**Figure 9.**
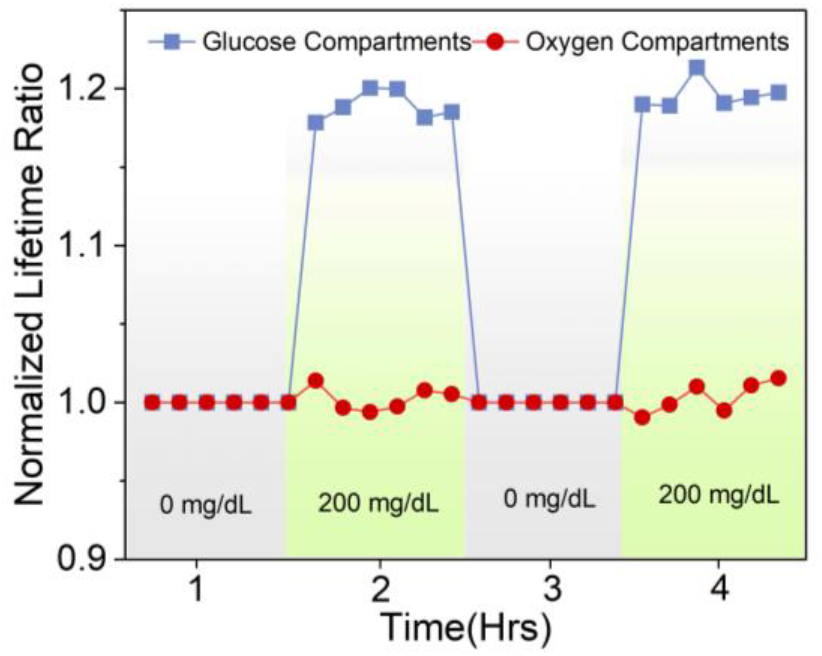
Phosphorescence lifetime (normalized to lifetime at 0 mg/dL glucose concentration) of barcode hydrogel sensor for glucose and oxygen-sensitive compartments under two cycles of 0 and 200 mg/dL glucose concentrations.

### *In Vivo* Studies – Histology

Histopathological evaluation provides the quantitative data needed to understand the long-term effects of insertable sensors in real-world conditions within the body. For this experiment, independent veterinary pathologists evaluated each sensor location and the surrounding tissue for device presence, device state, skin zone, host interface, and healing response (Figure 10). These sensor evaluations provide insight on the biocompatibility of the barcode sensors.

**Figure 10.**
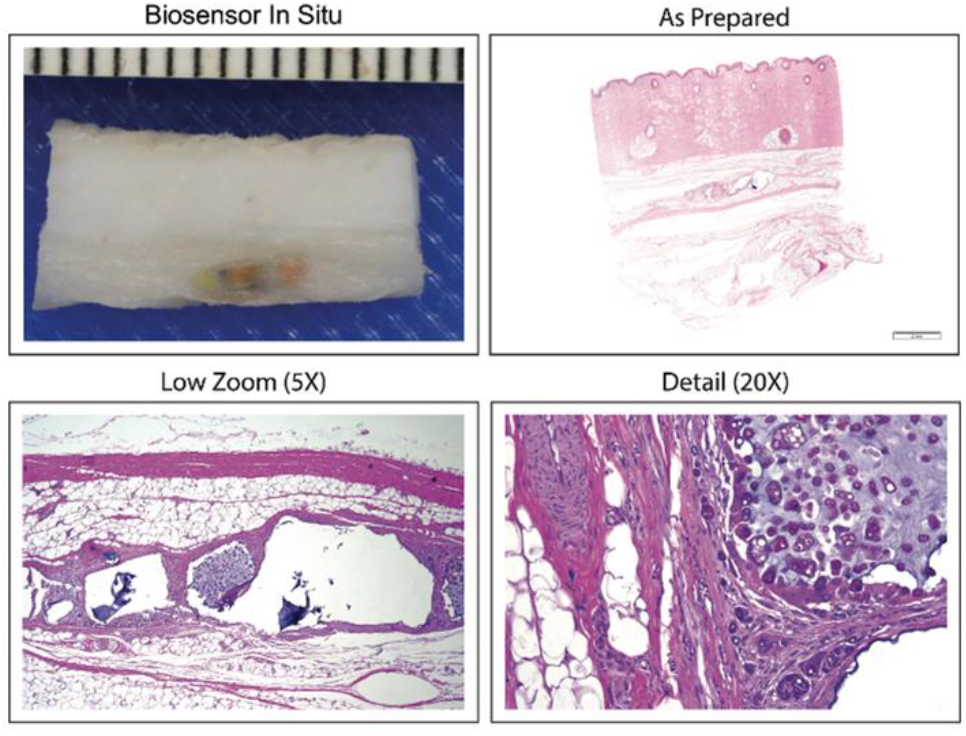
H&E-stained subcutaneous tissue section from porcine study with implanted barcode sensors. The sensors were well tolerated with minimal capsule formation and recovered intact and in good condition, even after 7 months in vivo.

Pig 1 was evaluated after the barcodes had been inserted for 7 months. (6/9) barcodes were still recoverable, and all barcodes that were located were externally readable and grossly visible on excision. The barcode sensors were mostly recovered from the subcutis (4/6) layers of the skin, with two of the eight sensors graded as being in the subcutis/muscle region (2/6). The depths from the surface of the skin to the shallowest portion of each device were also quantified, ranging from 3.84 to 9.19 mm deep, for an average depth of 6.05 mm.

In Pig 2, the barcodes were evaluated after 3 months of subcutaneous insertion. Every barcode could still be detected through the skin, and every barcode was grossly visible in the subcutaneous space after excision. Every barcode was found in the subcutis layer of the skin, with 2/7 barcodes graded as deep dermis/subcutis. The depth from the surface of the skin to the shallowest portion of each device ranged from 2.71 to 5.19 mm, for an average depth of 4.28 mm. The host interface of the barcodes was graded with mild (2/7) to moderate (5/7) fibrous capsule formation, and partial (4/7) to diffuse (2/7) infiltrative response from the surrounding tissue. On a single barcode, no infiltration was observed. The body’s healing response was graded as chronic phagocytosis (6/7), with one (1/7) barcode graded as chronic active phagocytosis. However, a subclinical sarcoptic mite population was detected on histology in Pig 2, which is likely responsible for the eosinophil seen on the slide. This conclusion and attribution of the deleterious response to the presence of mites is supported by the lack of chronic active response to the other sensors in either Pig 1 or 2, and in previous in vivo studies.

In both pigs, the sensors were primarily located in the subcutis. However, implants in the 7-month study in Pig 1 were graded as slightly deeper, erring on the muscle zone rather than the deep dermis zone. They were also somewhat (1.7mm) deeper in the skin. The different average depths seen in these studies could result from several factors. Firstly, implant positions were randomized, and pigs have widely varied natural skin thicknesses on various parts of their bodies. For example, sensors on the hind flank are naturally under much thinner skin than sensors under the tough shoulder region. Secondly, because pigs grow rapidly, the pigs doubled in size between the insertion procedures for Pigs 1 and 2, from about 60 pounds to around 120 pounds. This size difference changes the depth of the target subcutis zone. Finally, the sensors may change depth over time as skin thickens and fat deposits thicken as the pig matures. By the end of the experiment, the pigs weighed about 240 pounds each, and the pig’s skin grew along with the animals. The limited number of animals in this study precludes teasing apart these several factors. Fortunately, these modest depth variations did not affect the histological findings. More importantly, the ability to read implants at these depths demonstrates the effective penetration of the red excitation light and near-infrared emission through tissue (Figure S5).

In summary, the barcode sensors were well tolerated in healthy animals, remaining readable and visible in the subcutis at about 5mm under the skin. The typical host response was mild to moderate capsule formation, partial to diffuse cell infiltration, and chronic phagocytosis of the sensor material. Overall, this indicates that the devices are well tolerated over long-term subcutaneous insertion.

## CONCLUSION

In this study, we demonstrated a highly biocompatible insertable sensing platform for continuous multianalyte sensing, achieved by packaging different sensing assays within physically separated compartments within a molded hydrogel. The tunable assays capable of reporting oxygen and glucose at levels expected in tissue were realized by immobilizing oxygen-sensitive phosphors and oxidoreductase enzymes within alginate microparticles coated with polyelectrolyte nanofilms. The polyelectrolyte nanofilms provided the capability to precisely control diffusion into the discrete sensing microparticles to balance the diffusion-reaction kinetics at the physiologically relevant low oxygen concentrations. When subjected to cycles of different glucose concentrations, only the glucose-sensitive compartment demonstrated changes in phosphorescence lifetime while it remained constant in the oxygen-sensitive compartments, indicating no significant cross-interference between the compartments. Additionally, the barcode sensors demonstrated good stability for long-term storage, reproducibility, ability to withstand the sterilization process, and excellent biocompatibility. In the future, enzyme stabilization techniques will be exploited to prolong the lifespan of the barcode sensors. The addition of other analytes such as lactate, temperature, and uric acid will also be explored to broaden the applicability of the barcode sensors to other chronic conditions.

## ASSOCIATED CONTENT

### Supporting Information

Supplementary Table S1; Supplementary Figures S1-S4. Schematics of testing systems, Phosphorescence Lifetime response of oxygen barcodes, key metrics for barcode sensors with different sensing microparticles, raw data illustrating the change in phosphorescence lifetime over time with increases in oxygen concentration and glucose concentration, lifetime response for sensors incubated in glucose solution, signal-to-noise ratio for barcode sensors implanted in pigs (3months).

## AUTHOR INFORMATION

### Author Contributions

The manuscript was written through contributions of all authors. All authors have given approval to the version of the manuscript.

## ACKNOWLEDGMENT

This work was supported by the US National Science Foundation (NSF) Award # 2149551.

## REFERENCES

(1) Diabetes. World Health Organization, 2023. https://www.who.int/news-room/fact-sheets/detail/diabetes#:~:text=In%202019%2C%20diabetes%20was%20the,of%20cardiovascular%20deaths%20(1). (accessed 2023 December 30).

(2) Noncommunicable diseases. World health organization, 2023. (accessed 2024 01/23).

(3) de Meijer, V. E.; van’t Sant, H. P.; Spronk, S.; Kusters, F. J.; den Hoed, P. T. Reference value of transcutaneous oxygen measurement in diabetic patients compared with nondiabetic patients. Journal of Vascular Surgery 2008, 48 (2), 382–388. DOI: 10.1016/j.jvs.2008.03.010.

(4) Kanick, S. C.; Schneider, P. A.; Klitzman, B.; Wisniewski, N. A.; Rebrin, K. Continuous monitoring of interstitial tissue oxygen using subcutaneous oxygen microsensors: In vivo characterization in healthy volunteers. Microvascular Research 2019, 124, 6–18. DOI: 10.1016/j.mvr.2019.02.002.

(5) Das, S. K.; Nayak, K. K.; Krishnaswamy, P. R.; Kumar, V.; Bhat, N. Review—Electrochemistry and Other Emerging Technologies for Continuous Glucose Monitoring Devices. ECS Sensors Plus 2022, 1.

(6) Joseph, J. I. Review of the Long-Term Implantable Senseonics Continuous Glucose Monitoring System and Other Continuous Glucose Monitoring Systems. J Diabetes Sci Technol 2021, 15 (1), 167–173. From NLM.

(7) Jernelv, I. L.; Milenko, K.; Fuglerud, S. S.; Hjelme, D. R.; Ellingsen, R.; Aksnes, A. A review of optical methods for continuous glucose monitoring. Applied Spectroscopy Reviews 2019, 54 (7), 543–572. DOI: 10.1080/05704928.2018.1486324.

(8) Ahmed, I.; Jiang, N.; Shao, X.; Elsherif, M.; Alam, F.; Salih, A.; Butt, H.; Yetisen, A. K. Recent advances in optical sensors for continuous glucose monitoring. Sensors & Diagnostics 2022, 1 (6), 1098–1125, 10.1039/D1SD00030F. DOI: 10.1039/D1SD00030F.

(9) Wang, Y.; Vaddiraju, S.; Gu, B.; Papadimitrakopoulos, F.; Burgess, D. J. Foreign Body Reaction to Implantable Biosensors: Effects of Tissue Trauma and Implant Size. Journal of Diabetes Science and Technology 2015, 9 (5), 966–977. DOI: 10.1177/1932296815601869 (acccessed 2024/02/26).

(10) Nichols, S. P.; Koh, A.; Storm, W. L.; Shin, J. H.; Schoenfisch, M. H. Biocompatible materials for continuous glucose monitoring devices. Chem Rev 2013, 113 (4), 2528–2549. DOI: 10.1021/cr300387j From NLM.

(11) Tehrani, F.; Teymourian, H.; Wuerstle, B.; Kavner, J.; Patel, R.; Furmidge, A.; Aghavali, R.; Hosseini-Toudeshki, H.; Brown, C.; Zhang, F.; et al. An integrated wearable microneedle array for the continuous monitoring of multiple biomarkers in interstitial fluid. Nature Biomedical Engineering 2022, 6 (11), 1214–1224. DOI: 10.1038/s41551-022-00887-1.

(12) Glatz, R. T.; Ates, H. C.; Mohsenin, H.; Weber, W.; Dincer, C. Designing electrochemical microfluidic multiplexed biosensors for on-site applications. Analytical and Bioanalytical Chemistry 2022, 414 (22), 6531–6540. DOI: 10.1007/s00216-022-04210-4.

(13) Nichols, S. P.; Balaconis, M. K.; Gant, R. M.; Au-Yeung, K. Y.; Wisniewski, N. A. Long-Term In Vivo Oxygen Sensors for Peripheral Artery Disease Monitoring. In Oxygen Transport to Tissue XL, Thews, O., LaManna, J. C., Harrison, D. K. Eds.; Springer International Publishing, 2018; pp 351–356.

(14) Wisniewski, N. A.; Nichols, S. P.; Gamsey, S. J.; Pullins, S.; Au-Yeung, K. Y.; Klitzman, B.; Helton, K. L. Tissue-Integrating Oxygen Sensors: Continuous Tracking of Tissue Hypoxia. Adv Exp Med Biol 2017, 977, 377–383. DOI: 10.1007/978-3-319-55231-6_49 From NLM.

(15) Falcucci, T.; Presley, K. F.; Choi, J.; Fizpatrick, V.; Barry, J.; Kishore Sahoo, J.; Ly, J. T.; Grusenmeyer, T. A.; Dalton, M. J.; Kaplan, D. L. Degradable Silk-Based Subcutaneous Oxygen Sensors. Advanced Functional Materials 2022, 32 (27), 2202020. DOI: 10.1002/adfm.202202020.

(16) Andrus, L. P.; Unruh, R.; Wisniewski, N. A.; McShane, M. J. Characterization of Lactate Sensors Based on Lactate Oxidase and Palladium Benzoporphyrin Immobilized in Hydrogels. In Biosensors, 2015; Vol. 5, pp 398–416.

(17) Bornhoeft, L. R.; Biswas, A.; McShane, M. J. Composite Hydrogels with Engineered Microdomains for Optical Glucose Sensing at Low Oxygen Conditions. In Biosensors, 2017; Vol. 7.

(18) Falohun, T.; McShane, M. J. An Optical Urate Biosensor Based on Urate Oxidase and Long-Lifetime Metalloporphyrins. Sensors (Basel) 2020, 20 (4). DOI: 10.3390/s20040959 From NLM.

(19) Chen, Z.; Falohun, T.; Kameoka, J.; McShane, M. J. Multiplexed Implantable “Barcode” Hydrogel Platform for Continuous Oxygen and Glucose Monitoring. Texas A&M University 2024.

(20) Soundaram Jeevarathinam, A.; Saleem, W.; Martin, N.; Hu, C.; McShane, M. J. NIR Luminescent Oxygen-Sensing Nanoparticles for Continuous Glucose and Lactate Monitoring. In Biosensors, 2023; Vol. 13.

(21) Ko, B.; Zavareh, A. T.; McShane, M. J. In Vitro System for Evaluation of Biosensors in Controlled Dynamic Environmental Conditions. IEEE Transactions on Instrumentation and Measurement 2024, 73, 1–12. DOI: 10.1109/TIM.2023.3334366.

(22) Zavareh, A. T.; Ko, B.; Roberts, J.; Elahi, S.; Mcshane, M. J. A Versatile Multichannel Instrument for Measurement of Ratiometric Fluorescence Intensity and Phosphorescence Lifetime. IEEE Access 2021, 9, 103835–103849. DOI: 10.1109/ACCESS.2021.3098777.

(23) Goncharov, A.; Gorocs, Z.; Pradhan, R.; Ko, B.; Ajmal, A.; Rodriguez, A.; Baum, D.; Veszpremi, M.; Yang, X.; Pindrys, M. An insertable glucose sensor using a compact and cost-effective phosphorescence lifetime imager and machine learning. arXiv preprint arXiv:2406.09442 2024.

(24) Enzymatic Assay of Glucose Oxidase. https://www.sigmaaldrich.com/US/en/technical-documents/protocol/protein-biology/enzyme-activity-assays/enzymatic-assay-of-glucose-oxidase (accessed 03/18/2023).

(25) Zhu, H.; Srivastava, R.; Brown, J. Q.; McShane, M. J. Combined physical and chemical immobilization of glucose oxidase in alginate microspheres improves stability of encapsulation and activity. Bioconjug Chem 2005, 16 (6), 1451–1458. DOI: 10.1021/bc050171z From NLM.

(26) Biswas, A.; Nagaraja, A. T.; You, Y.-H.; Roberts, J. R.; McShane, M. J. Cross-linked nanofilms for tunable permeability control in a composite microdomain system. RSC Advances 2016, 6 (75), 71781–71790, 10.1039/C6RA13507B. DOI: 10.1039/C6RA13507B.

(27) von Woedtke, T.; Jülich, W. D.; Hartmann, V.; Stieber, M.; Abel, P. U. Sterilization of enzyme glucose sensors: problems and concepts. Biosensors and Bioelectronics 2002, 17 (5), 373–382. DOI: 10.1016/S0956-5663(01)00310-4.

(28) Fuchs, S.; Rieger, V.; Tjell, A. Ø.; Spitz, S.; Brandauer, K.; Schaller-Ammann, R.; Feiel, J.; Ertl, P.; Klimant, I.; Mayr, T. Optical glucose sensor for microfluidic cell culture systems. Biosensors and Bioelectronics 2023, 237, 115491. DOI: 10.1016/j.bios.2023.115491.

(29) Mross, S.; Zimmermann, T.; Zenzes, S.; Kraft, M.; Vogt, H. Study of enzyme sensors with wide, adjustable measurement ranges for in-situ monitoring of biotechnological processes. Sensors and Actuators B: Chemical 2017, 241, 48–54. DOI: 10.1016/j.snb.2016.10.054.

(30) Dang, T. T.; Aroyan, S.; Keistensen, J. S. Protective agents against e-beam irradiation for proteins in optical sensing chemistry. US 2015.

(31) Suvarli, N.; Wenger, L.; Serra, C.; Perner-Nochta, I.; Hubbuch, J.; Wörner, M. Immobilization of β-Galactosidase by Encapsulation of Enzyme-Conjugated Polymer Nanoparticles Inside Hydrogel Microparticles. Front Bioeng Biotechnol 2021, 9, 818053. DOI: 10.3389/fbioe.2021.818053 From NLM.

(32) Park, J.; Kim, J.; Kim, S.-Y.; Cheong, W. H.; Jang, J.; Park, Y.-G.; Na, K.; Kim, Y.-T.; Heo, J. H.; Lee, C. Y.; et al. Soft, smart contact lenses with integrations of wireless circuits, glucose sensors, and displays. Science Advances 4 (1), eaap9841. DOI: 10.1126/sciadv.aap9841.

