## Supporting Information for "Continuous Monitoring of Glucose and Oxygen using an Insertable Biomaterial-based Multianalyte Barcode Sensor"

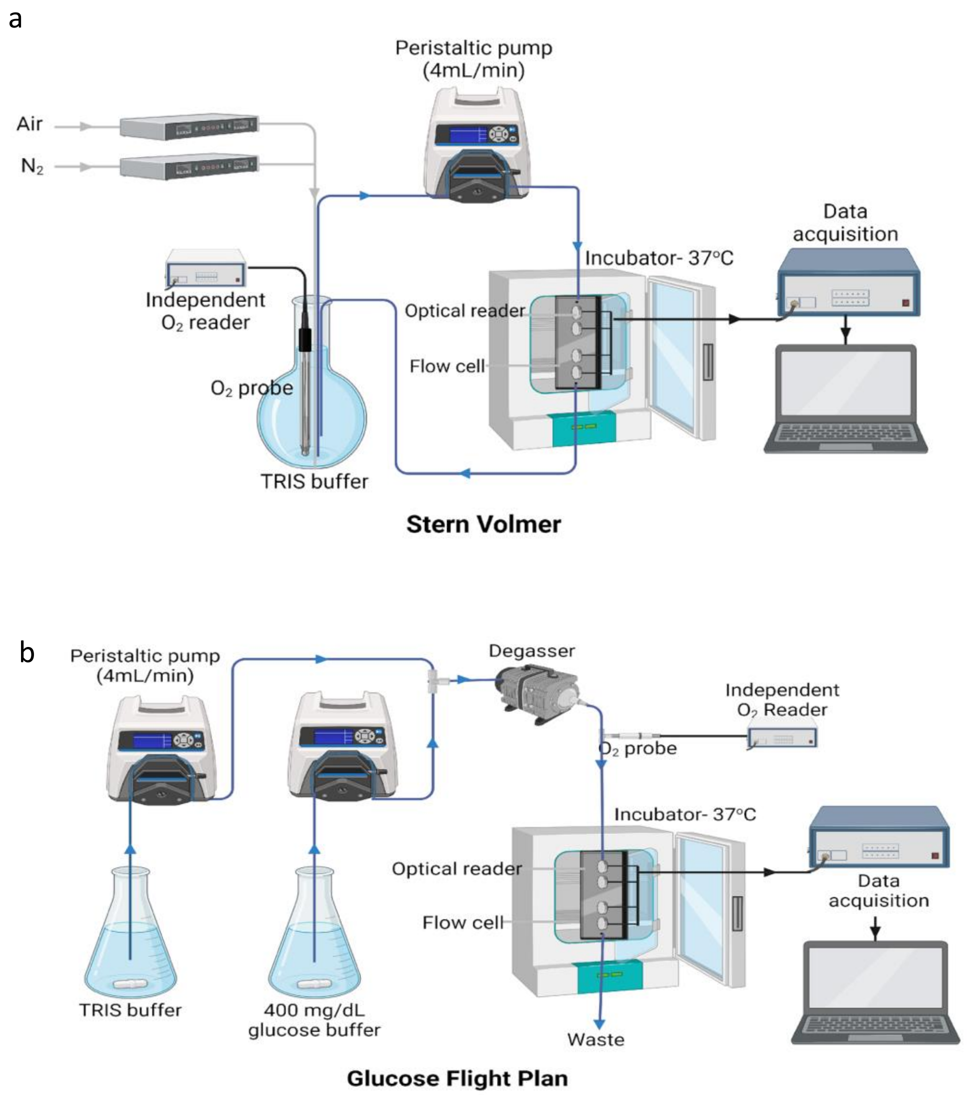


**Figure S1**: Schematic representation of **(a)**oxygen response testing apparatus, and **(b)** glucose flight plan testing system.

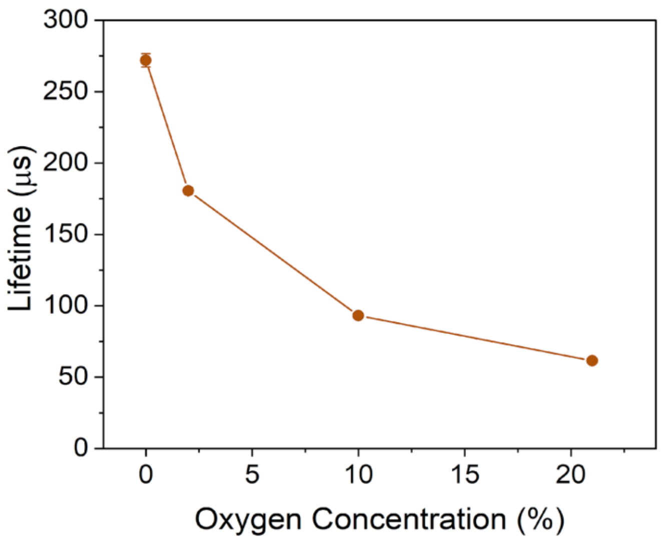


**Figure S2:** (a) Phosphorescence lifetime response oxygen barcode hydrogel sensor under different oxygen concentrations. Error bars represent the SD from the mean for N=4 different samples.

| | LOD[a] (mg/dL) | MDGC[b] (mg/dL) | Range[c] (mg/dL) | Sensitivity (µs-dL/mg) | Oxygen Sensitivity ($K_{SV}$, $\%O_2^{-1}$)[d] |
|---|---|---|---|---|---|
| **Non-crosslinked** | | | | | |
| 5 LbL | 10.7 | 65.0 | 54.3 | 0.87 | 0.16 |
| 10 LbL | 15.3 | 77.2 | 61.9 | 0.72 | 0.14 |
| 15 LbL | 14.0 | 109.0 | 95.1 | 0.50 | 0.14 |
| 20 LbL | 11.2 | 112.4 | 101.2 | 0.52 | 0.14 |
| 25 LbL | 14.6 | 147.8 | 133.2 | 0.37 | 0.16 |
| **Crosslinked** | | | | | |
| 25 CXL | 21.3 | 175.5 | 154.3 | 0.38 | 0.15 |
| 30 CXL | 37.8 | 221.3 | 183.5 | 0.20 | 0.15 |
| 35 CXL | 37.1 | 321.1 | 284.0 | 0.19 | 0.15 |

**Table S1:** Figures of merit calculated for barcode hydrogels with microparticles having uncrosslinked nanofilm coatings (LbL) and crosslinked (CXL) nanofilm coatings comprising different numbers of polyelectrolyte bilayers. Each data represents the average of measurement from 4 different samples. [a]LOD, Limit of detection. [b]MDGC, maximum differentiable glucose concentration. [c]Range, MDGC- LOD. [d]$K_{SV}$, Stern-Volmer constant.

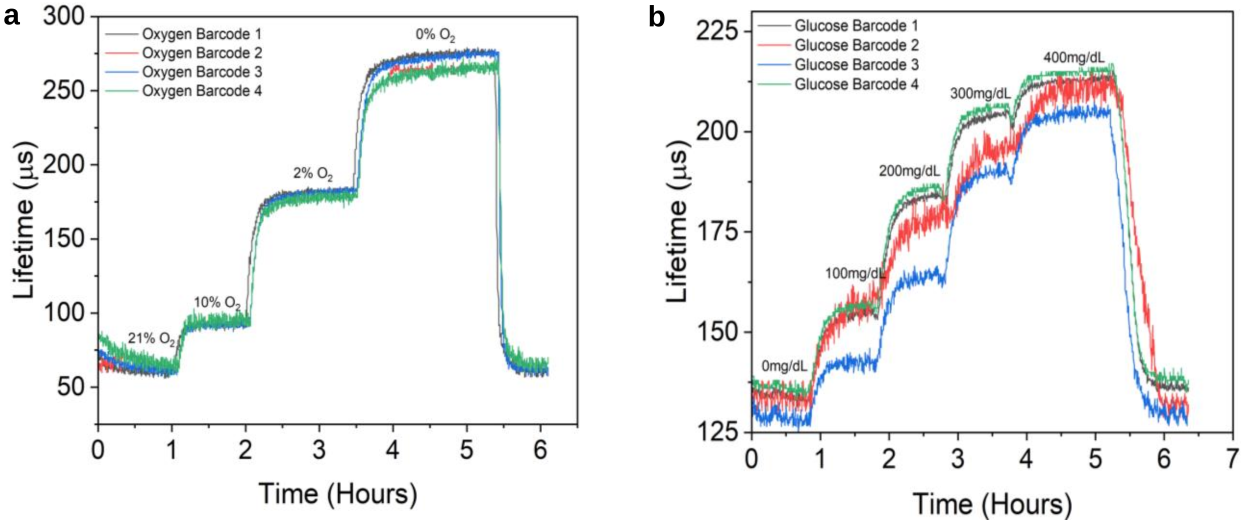


**Figure S3: (a)** Phosphorescence lifetime response over time to changing dissolved oxygen concentrations for oxygen barcodes. **(b)** Phosphorescence lifetime response over time to changing glucose concentration at fixed oxygen (~30µM) for glucose barcodes. The average and standard deviations of lifetime measurements were calculated from the plateaus for these data, which were then presented as the steady-state lifetime responses in other graphs (Figures 4, 5, 6, 7, 8a, 9, and S4)

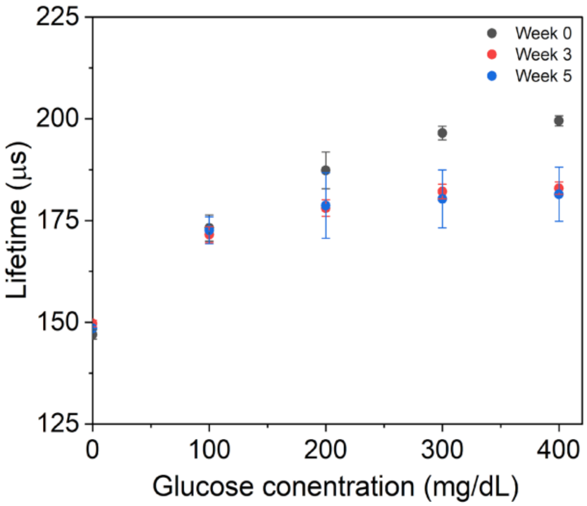


**Figure S4**: Lifetime response of glucose barcodes stored in PBS (pH 7.4) with 100 mg/dL of glucose at 37º C at time intervals of weeks 0, 3, and 5. Error bars represent the SD of the mean for N=4 different samples.

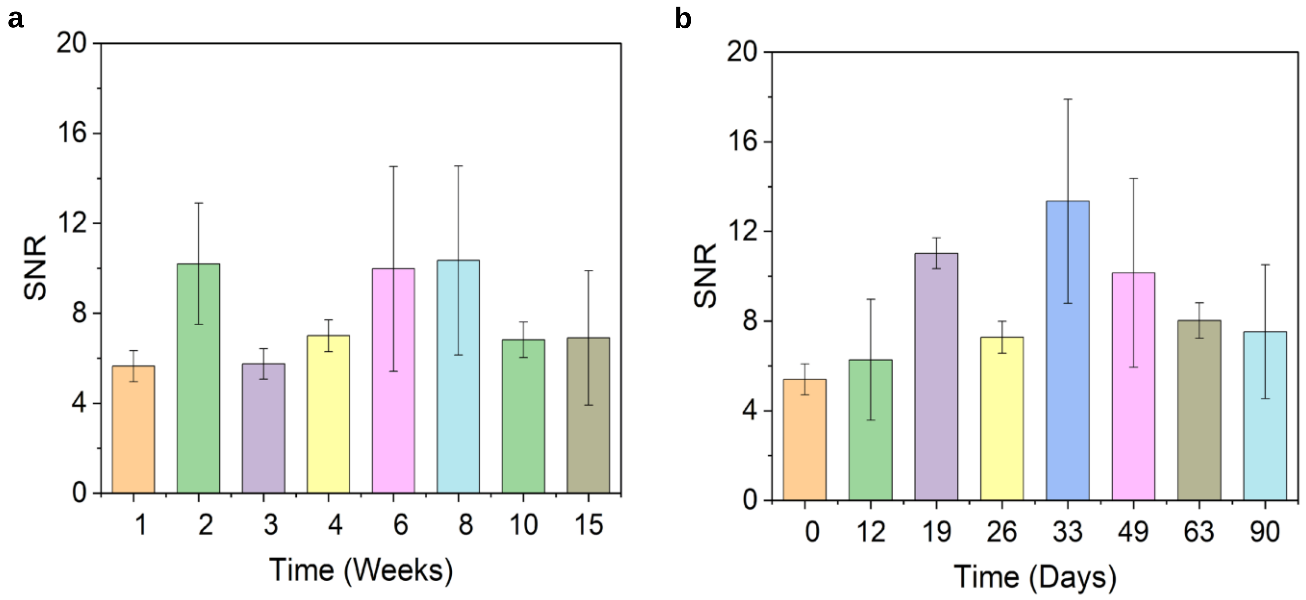


**Figure S5**: Signal-to-noise ratio (SNR) of barcode sensors implanted in (a) Pig 1 and (b) Pig 2 for a duration of over 3 months. Error bars represent the SD from the mean for N=4 different samples.
